# The rumen microbiome: an underexplored resource for novel antimicrobial discovery

**DOI:** 10.1101/148221

**Authors:** Linda B Oyama, Susan E Girdwood, Alan R Cookson, Narcis Fernandez-Fuentes, Florence Privé, Hannah E Vallin, Toby J Wilkinson, Peter N Golyshin, Olga V Golyshina, Ralf Mikut, Kai Hilpert, Jennifer Richards, Mandy Wootton, Joan E Edwards, Marc Maresca, Josette Perrier, Fionnuala T Lundy, Yu Luo, Mei Zhou, Matthias Hess, Hilario C Mantovani, Christopher J Creevey, Sharon A Huws

## Abstract

Antimicrobial peptides (AMPs) are promising drug candidates to target multi-drug resistant bacteria. The rumen microbiome presents an underexplored resource for the discovery of novel microbial enzymes and metabolites, including AMPs. Using functional screening and computational approaches, we identified 181 potentially novel AMPs from a rumen bacterial metagenome. Here, we show that three of the selected AMPs (Lynronne-1, 2 and 3) were effective against numerous bacterial pathogens, including methicillin resistant *Staphylococcus aureus* (MRSA). No decrease in MRSA susceptibility was observed after 25 days of sub-lethal exposure to these AMPs. The AMPs bound preferentially to bacterial membrane lipids and induced membrane permeability leading to cytoplasmic leakage. Topical administration of Lynronne-1 (10% w/v) to a mouse model of MRSA wound infection elicited a significant reduction in bacterial counts, which was comparable to treatment with 2% mupirocin ointment. Our findings indicate that the rumen microbiome may provide viable alternative antimicrobials for future therapeutic application.

## Introduction

The global increase of multi-drug resistant (MDR) bacteria, combined with the decreasing number of novel and effective antibacterial agents being developed for the market, poses a serious global threat to human health ^1,2^. Urgency is required with respect to discovery and development of novel antibiotics, alongside antimicrobial stewardship and development of rapid tools to detect MDR bacteria ^3^. Antimicrobial peptides (AMPs), particularly cationic AMPs, are promising alternatives to currently available antibiotics, due to their broad spectrum activity against Gram positive and Gram negative bacteria, and their rapid mode of action on the cytoplasmic membrane, which results in a decreased rate of resistance being developed by the targeted pathogens. However, synthetic generation of AMPs has produced very few if any viable AMPs for medical use, thus there is a need to further prospect from nature ^4^.

Bacterial communities are often symbiotic, but on occasion may need to compete for resources to survive. In order to compete, bacteria often up-regulate genes encoding AMPs to kill other bacteria in the community ^5^. The rumen is an example of such a highly competitive community with a very complex microbiome, composed of unique and diverse range of bacteria, archaea, protozoa, fungi, and viruses ^6,7^ and as such is a potentially unique resource for novel AMP discovery ^8-10^. In the past few years, it has become apparent that culture dependent methods to evaluate microbial populations substantially underestimates the diversity of microorganisms present, and only around 10-20% of the species from the rumen environment have been cultured to date ^11,12^. The limitations surrounding culturability and antimicrobial discovery can be circumvented using sequence and functional based metagenomics, which allow mining of whole microbiomes for novel bioactive compounds ^13,14^

Here, we report the discovery and characterization of three AMPs (i.e. Lynronne-1, Lynronne-2 and Lynronne-3) via functional metagenomic screening of a rumen bacterial metagenomic library and *in-silico* sequence data analysis. These novel AMPs from the rumen microbiome have the potential to treat MDR bacterial infections.

## Results

### Identification of antimicrobial peptides

A metagenomic library consisting of 8,448 fosmids from the plant-attached rumen microbiome ^15^ was screened for antimicrobial activity by a combination of agar-based functional screening, sequencing of positive clones and bioinformatic prediction of AMP sequences. To test the antimicrobial activity of fosmid clones, lawns of pathogens including methicillin sensitive *Staphylococcus aureus* (MSSA) RN4220, *Escherichia coli* K12, *Salmonella enterica* serovar Typhimurium SL1344, *Listeria monocytogenes* NCTC 11994 (serovar 4b) and *Enterococcus faecalis* JH2-2 were inoculated on appropriate agar medium and the clones were gently transferred on top using multichannel pin replicators. After an incubation period of 24 h at the appropriate temperatures, 255 fosmids produced clearing zones in the pathogen lawn, indicating that the metagenome clones carried inserts with antimicrobial activity. Of these clones, twenty-four were selected randomly and sequenced using the GS FLX system. A summary of the assembly metrics of the twenty-four metagenomic inserts with antimicrobial activity is provided in Extended Data Table 1. The full fosmid sequences are available in the GenBank database under the BioProject PRJNA264884; BioSample IDs: SAMN063330279 to SAMN063330302. Prediction of AMPs from the identified ORFs yielded a total of 181 AMPs. A peptide library consisting of 135 of these AMPs (≤25 AAs) was synthesized and screened for antimicrobial activity using the SPOT technique ^16^. Twenty-five active AMPs were identified in this screen (Extended Data Table 2). Candidates, Lynronne-1 (19 AAs: LPRRNRWSKIWKKVVTVFS-NH_2_), Lynronne-2 (20 AAs: HLRRINKLLTRIGLYRHAFG-NH_2_) and Lynronne-3 (20 AAs: NRFTARFRRTPWRLCLQFRQ-NH_2_) encoded in fosmid clones SABPL29H11, SABPL5A1 and SABPL12(2)A3 respectively and were selected for further analysis due to their broad spectrum antibacterial activity and lower MIC values. The nucleotide and protein sequences of the ORFs from which they were derived are available in the GenBank database under accession numbers KY628802, KY628803 and KY628804, respectively. Sequences upstream and downstream of these ORFs are shown in Extended Data Fig. 2. Structural modelling using PEP-FOLD ^17^ indicate that these peptides adopt a α-helical conformation of amphipathic nature, an arrangement typical of many α-helical antimicrobial peptides^18^ (Fig. 1). The AMPs have a net positive charge of +6, +5 and +6, respectively with a hydrophobicity ratio of ≥40%.

**Fig. 1.**
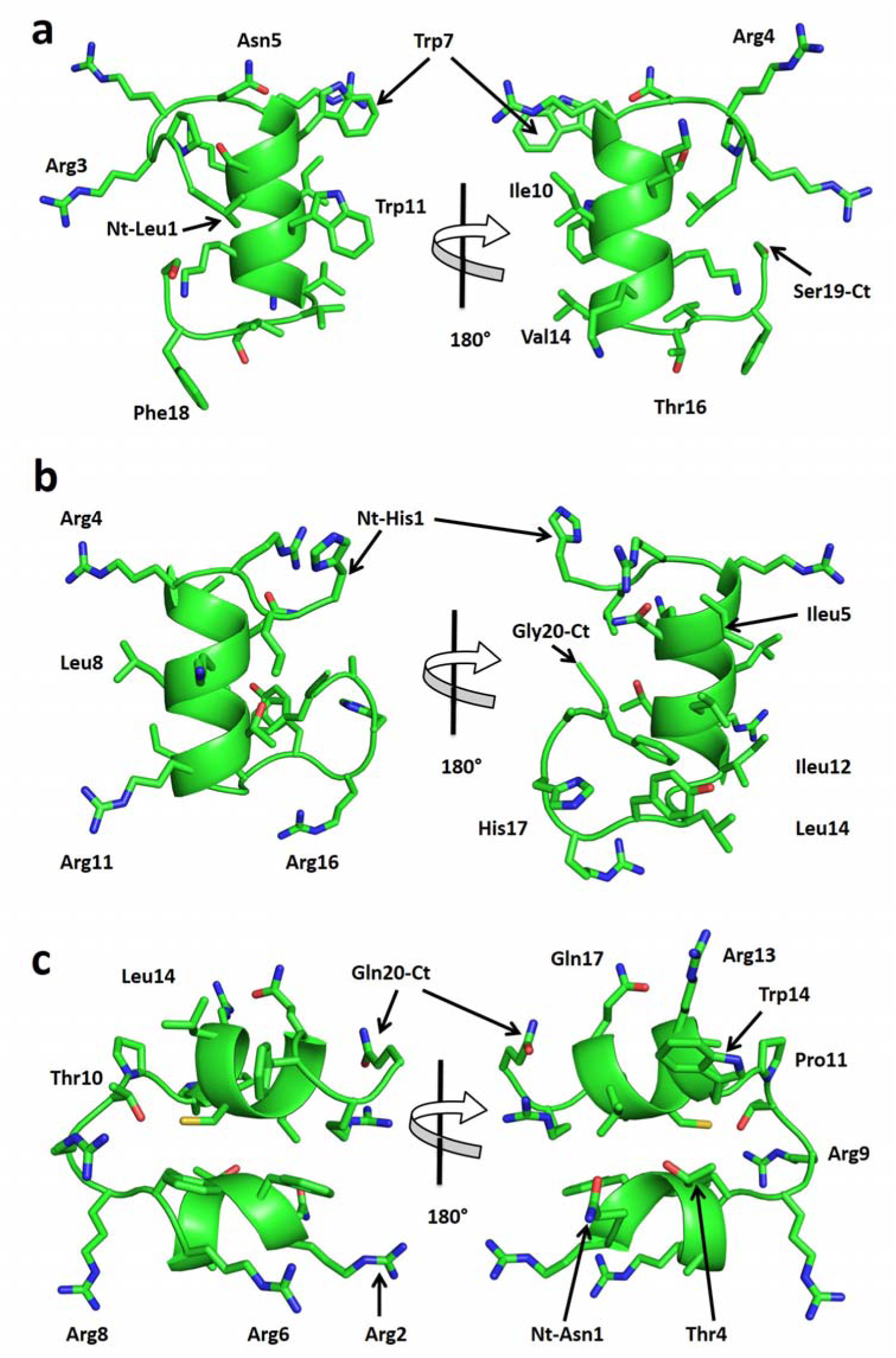
Predicted structures for peptides: Lynronne-1, 2 and 3. **a)** Lynronne-1, **b)** Lynronne-2 and **c)** Lynronne-3. Main-chain and side chains depicted in ribbon and stick representation respectively and colored according to atom type: Carbon, Oxygen and Nitrogen in green, red and blue respective. Two orientations are shown rotated about the shown axis. Ct and Nt as well as selected residues are depicted in the figure. Figures were rendered using PyMol.

**Table 1:** **Minimum Inhibitory Concentrations (MICs) against drug resistant pathogens.**

| Organism information |  |  | Peptides and comparator antibiotics (μg/ml) |  |  |  |  |  |  |  |  |  |  |  |  |  |
| --- | --- | --- | --- | --- | --- | --- | --- | --- | --- | --- | --- | --- | --- | --- | --- | --- |
| Lab no./Strain ID | Organism | Resistances | Cip/<br>Lev (L) | L-1 | L-2 | L-3 | P4 | P5 | P8 | P10 | P11 | P12 | P15a | P15s | Van | Mup |
| EMRSA-15 | *S. aureus* | MRSA, Cip | >256 | 32 | 32 | 32 | 128 | 64 | 64 | 250 | 250 | 250 |  | 128 | 2 |  |
| ATCC 33591 | *S. aureus* | MRSA |  | 16 | 128 | 64 |  |  |  |  |  |  |  |  | 2 |  |
| USA300 BAA-1717 | *S. aureus* | MRSA |  | 8 | 256 | 64 |  |  |  |  |  |  |  |  | 2 | 0.12 |
| Q14-0320 | *S. aureus* | MRSA | 64 | 32 | >128 | 128 | >128 | >128 | >128 | >128 | 64 | 64 | >128 | >128 |  |  |
| Q14-0325 | *S. aureus* | MRSA | 0.25 | 32 | >128 | 128 | >128 | 128 | 128 | >128 | 32 | 64 | >128 | 64 |  |  |
| Q14-0352 | *S. aureus* | MRSA | 0.5 | 32 | >128 | 128 | >128 | >128 | 128 | >128 | 32 | 64 | >128 | >128 |  |  |
| Q14-0390 | *S. aureus* | MRSA | 64 | 32 | >128 | 128 | >128 | 128 | 128 | >128 | 32 | 64 | >128 | >128 |  |  |
| NCTC 12493 | *S. aureus* | MRSA | 0.25 | 16 | 64 | 64 | 32 | 64 | 32 | >128 | 32 | 16 | >128 | 64 |  |  |
| ATCC 29213 | *S. aureus* | Sensitive | 0.25 | 32 | >128 | 128 | >128 | 128 | 128 | >128 | 32 | 64 | >128 | >128 |  |  |
| RN4220 | *S. aureus* | Sensitive | >256 | 32 | 32 | 32 | 128 | 64 | 64 | 256 | 256 | 256 |  | 128 | 1 |  |
| Q14-0095 | *K. pneumoniae* | KPC | 16 | 32 | >128 | 32 | >128 | 128 | >128 | >128 | >128 | >128 | >128 | 128 |  |  |
| Q14-0146 | *K. pneumoniae* | OXA-48 | >128 | 32 | 64 | 64 | >128 | >128 | >128 | >128 | >128 | >128 | >128 | 128 |  |  |
| Q14-0285 | *K. pneumoniae* | VIM | 32 | 16 | 64 | 64 | 64 | 128 | 128 | >128 | >128 | >128 | >128 | 128 |  |  |
| Q14-0251 | *K. pneumoniae* | CTX-M | 16 | 16 | 64 | 64 | >128 | >128 | >128 | >128 | 128 | 128 | >128 | 128 |  |  |
| ATCC 700603 | *K. pneumoniae* | SHV-18 | 0.25 | 16 | 64 | 32 | 128 | >128 | 128 | >128 | 128 | 64 | >128 | 128 |  |  |
| NCTC 13442 | *K. pneumoniae* | OXA-48 | 1 | 16 | 128 | 32 | >128 | 128 | >128 | >128 | 128 | >128 | >128 | 128 |  |  |
|  | *K. pneumoniae* | Sensitive | 0.03 | 16 | 128 | 32 | >128 | >128 | >128 | >128 | 128 | >128 | >128 | >128 |  |  |
| Q13-0717 | *A. baumannii* | IMI, MER | 16 | 4 | 8 | 32 | >128 | 32 | 64 | >128 | 64 | 64 | >128 | 128 |  |  |
| Q13-0795 | *A. baumannii* | IMI, MER | 16 | 4 | 8 | 32 | >128 | 32 | 64 | >128 | 64 | 64 | 128 | 64 |  |  |
| 1560739339 | *A. baumannii* | OXA-23 | 128 | 4 | 16 | 32 | >128 | 128 | 64 | >128 | 128 | 64 | 128 | 128 |  |  |
| Q12-0809 | *A. baumannii* | OXA-23, OXA-51 | 128 | 4 | 8 | 16 | 32 | 64 | 32 | 64 | 128 | 32 | 128 | 128 |  |  |
| Q12-0867 | *A. baumannii* | Sensitive | 0.25 | 4 | 8 | 8 | 64 | 16 | 64 | 128 | 32 | 16 | 128 | 64 |  |  |
| Q14-0208 | *P. aeruginosa* (CF) | CAZ, IMI, MER | 4 | 16 | 32 | 32 | >128 | 128 | 64 | >128 | 64 | 64 | 128 | 128 |  |  |
| Q14-0890 | *P. aeruginosa* (CF) | CAZ, IMI, MER | 4 | 32 | 64 | 16 | 64 | 64 | 32 | 128 | 16 | 32 | 128 | 64 |  |  |
| Q12-0535 | *P. aeruginosa* (CF) | Sensitive | 0.06 | 64 | 128 | 128 | >128 | 128 | 128 | 125 | 128 | 32 | 128 | 128 |  |  |
| Q12-0581 | *P. aeruginosa* (CF) | IMI, MER | 2 | 32 | 128 | 32 | 128 | 64 | 128 | >128 | 32 | 128 | 128 | 64 |  |  |
| Q12-0758 | *P. aeruginosa* (CF) | CAZ, IMI, MER | 2 | 32 | 64 | 32 | 128 | 128 | 128 | >128 | 32 | 128 | 128 | 64 |  |  |
| ATCC 27853 | *P. aeruginosa* (CF) | Sensitive | 0.25 | 64 | >128 | 128 | >128 | 128 | >128 | >128 | >128 | 128 | 128 | 128 |  |  |
| PA01 | *P. aeruginosa* (CF) |  | 0.5 | 32 | 128 | 32 | 512 | 64 | 32 | 512 | 512 | 512 |  | 128 |  |  |
| AMT0060 | *P. aeruginosa* (CF) |  | 0.06 (L) | 32 | 64 | 128 |  |  |  |  |  |  | >512 | 64 |  |  |
| C3719 | *P. aeruginosa* (CF) |  | <0.03 (L) | 64 | 32 | 16 |  |  |  |  |  |  | >512 | 32 |  |  |
| LESB58 | *P. aeruginosa* (CF) |  | 0.06 (L) | 64 | 256 | 32 |  |  |  |  |  |  | >512 | 128 |  |  |
| M15U114296A | *C. coli* | Cip, Tet, Ery, Amox, Gent | >128 | 64 | >128 | 128 | >128 | 128 | >128 | >128 | >128 | >128 | 128 | 128 |  |  |
| M15U114452A | *C. coli* | Cip, Cla, Ery | >128 | 64 | >128 | 128 | >128 | 128 | >128 | >128 | >128 | >128 | 128 | 128 |  |  |
| M16U109179A | *C. jejuni* | Azith, Cip, Ery, Tet | >128 | 64 | >128 | 128 | >128 | 128 | >128 | >128 | >128 | >128 | 128 | 128 |  |  |
| M16U109568A | *C. jejuni* | Cip | 128 | 64 | >128 | 64 | >128 | 128 | >128 | >128 | >128 | >128 | 128 | 128 |  |  |
| NCTC 11351 | *C. jejuni* |  |  | 64 | >128 | 128 | >128 | 128 | >128 | >128 | >128 | >128 | 128 | 128 |  |  |
| NCTC 11161 | *C. jejuni* |  | 0.5 | 4 | 32 | 256 |  |  |  |  |  |  |  |  |  |  |
| ATCC 33292 | *C. jejuni* |  | 0.12 | 4 | 32 | 64 |  |  |  |  |  |  |  |  |  |  |
| K12 | *E. coli* |  | 0.06 | 32 | 64 | 64 | 64 | 64 | 64 | 256 | 256 | 256 |  | 64 | 128 |  |
| SL1344 | *Sal. typhimurium* |  | 0.12 | 32 | 32 | 64 | 32 | 64 | 32 | 64 | 512 | 256 |  | 128 | 256 |  |
| JH2-2 | *Ent. faecalis* |  | 64 | 32 | 32 | 64 | 64 | 64 | 64 | 512 | 256 | 256 |  | 128 | 64 |  |
| NCTC 11994 | *List. monocytogenes* |  | 64 | 32 | 32 | 32 | 64 | 64 | 64 | 512 | 256 | 128 |  | 128 | 64 |  |

L-1, L-2 and L-3: Lynronne-1, 2 and 3, P4-P15a: Peptides 4 to 15a, Lev (L): Levofloxacin, Cip: Ciprofloxacin, Van: Vancomycin, Mup; Mupirocin, KPC: *Klebsiella pneumoniae* carbapenemase, OXA: Oxacillin, VIM: Metallo- $\beta$ -Lactamase gene, CTX-M: extended spectrum $\beta$ -Lactamase, SHV-18: $\beta$ -Lactamase, IMI (imipenem): MER (meropenem) CAZ (ceftazidime), Tet: Tetracycline, Ery: Erythromycin, Amox: Amoxicillin, Gent: Gentamicin, Azith: Azithromycin, Cla: Clarithromycin. Blank: Not tested.

### Antimicrobial activity, resistance and mechanism of action

The antibacterial activity of Lynronne-1, Lynronne-2 and Lynronne-3 against twenty-nine clinically important multidrug-resistant pathogens was explored. All three peptides had promising activity against Gram-positive pathogens, including MDR strains (Table 1). Lynronne-1 had a minimum inhibitory concentration (MIC) range of 8-32, Lynronne-2: 32-256 and Lynronne-3: 32-128 of *μ*g/ml for MRSA strains. Since cationic peptides rarely demonstrate high *in vitro* antimicrobial activities seen with some conventional antibiotics, the MICs determined for Lynronne-1, 2 and 3 were not surprising and fall within the range of many commercially available AMPs ^19,20^.

Lynronne-1, 2 and 3 (at 3x MIC) had relatively faster bactericidal activity against three MRSA strains when compared to vancomycin, a first line drug for MRSA (Fig. 2a, and Extended Data Fig 3a, b). Reductions of >8 log_10_ CFU/ml occurred within the first 10 mins for Lynronne-1 and Lynronne-3, and >3 log_10_ CFU/ml in 30 mins for Lynronne-2 (Fig. 2a, and Extended Data Fig 3a, b). Vancomycin at 3x MIC produced between 1 to >4 log_10_ CFU/ml reductions depending on the MRSA strain. This can be attributed to difference in kill kinetics and mode of action ^21^. Time dependent killing of *Escherichia coli* K12 was investigated to ascertain the efficacy of the peptides against Gram-negative bacterial strains. All three AMPs caused a rapid decrease in *E. coli* K12 viability at 3x MIC with complete cell death occurring within 10 mins (see Extended Data Fig. 1a). Control agents, Polymyxin B and ciprofloxacin showed expected bactericidal activities ^22^ (>4 log_10_ CFU/ml) against *E. coli* K12 at 10 mins and 3 h respectively (see Extended Data Fig. 1a).

**Fig. 2.**
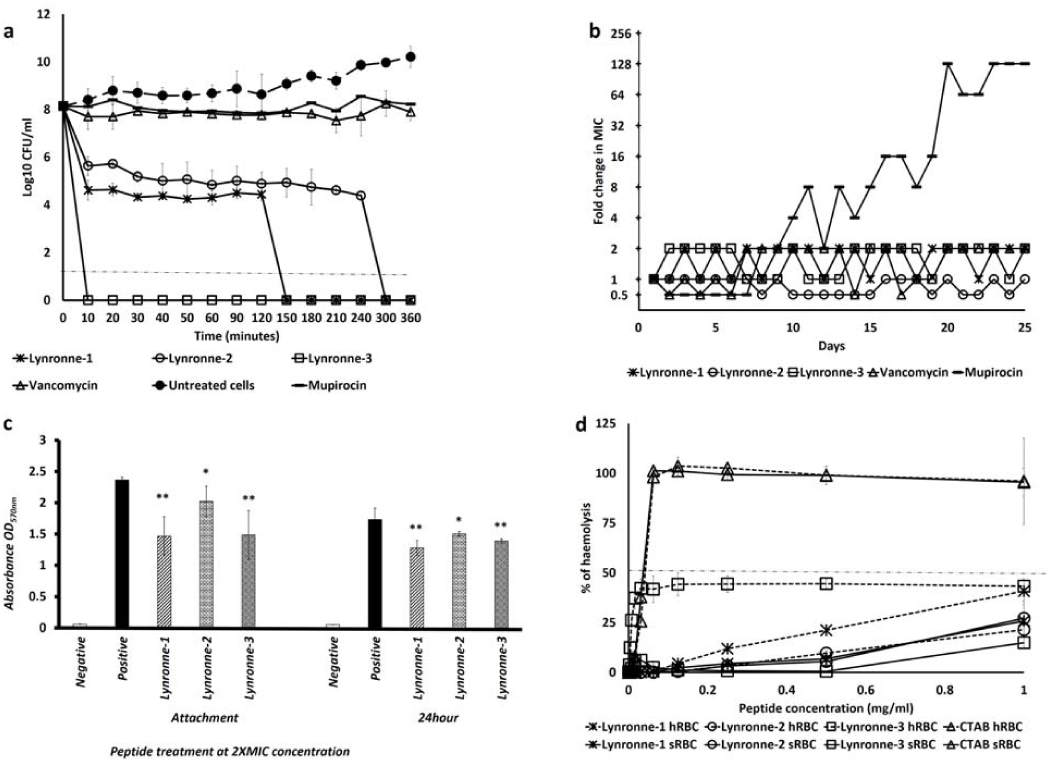
Antimicrobial activity and haemolytic activity of Lynronne-1, 2 and 3. **a)** Time dependent kill of MRSA USA300 by antimicrobial compounds at 3x MIC concentration. Dashed lines indicate limit of detection. **b)** Resistance acquisition during serial passaging of MRSA USA300 cells in the presence of sub-MIC levels of antimicrobials. The y axis is the fold change in MIC during passaging. For mupirocin, 256x MIC was the highest concentration tested. The figure is representative of 3 independent experiments. **c)** Quantification of anti-biofilm activity (MRSA 33591): * and ** (P < 0.05 and 0.01 respectively-significantly different from untreated cells (positive). **d)** Haemolytic activity, dashed line indicates 50% haemolysis. Values from at least three independent replicates; error bars represent the standard deviation).

Serial passage of MRSA strains in the presence of sub-MIC levels of Lynronne-1, 2 and 3 over a period of 25 days failed to produce mutants that were resistant to the AMPs (Fig. 2b and Extended Data Fig. 3c, d), suggesting a non-specific mode of action ^23^. The efficacy of Lynronne-1, 2 and 3 against *S. aureus* biofilm attachment and established biofilms at sub and supra-MIC concentrations were tested. MRSA ATCC 33591 strain was used in this assay as it was a better biofilm former in our hands than the other strains (results not shown). All AMPs decreased biofilm attachment at 2x MIC concentrations (Fig. 2c) and Lynronne-2 was effective even at 0.25x MIC (data not shown). Anti-biofilm activity against established biofilms was observed with all peptides at ≥ 2x MIC concentrations (Fig. 2c).

**Fig. 3.**
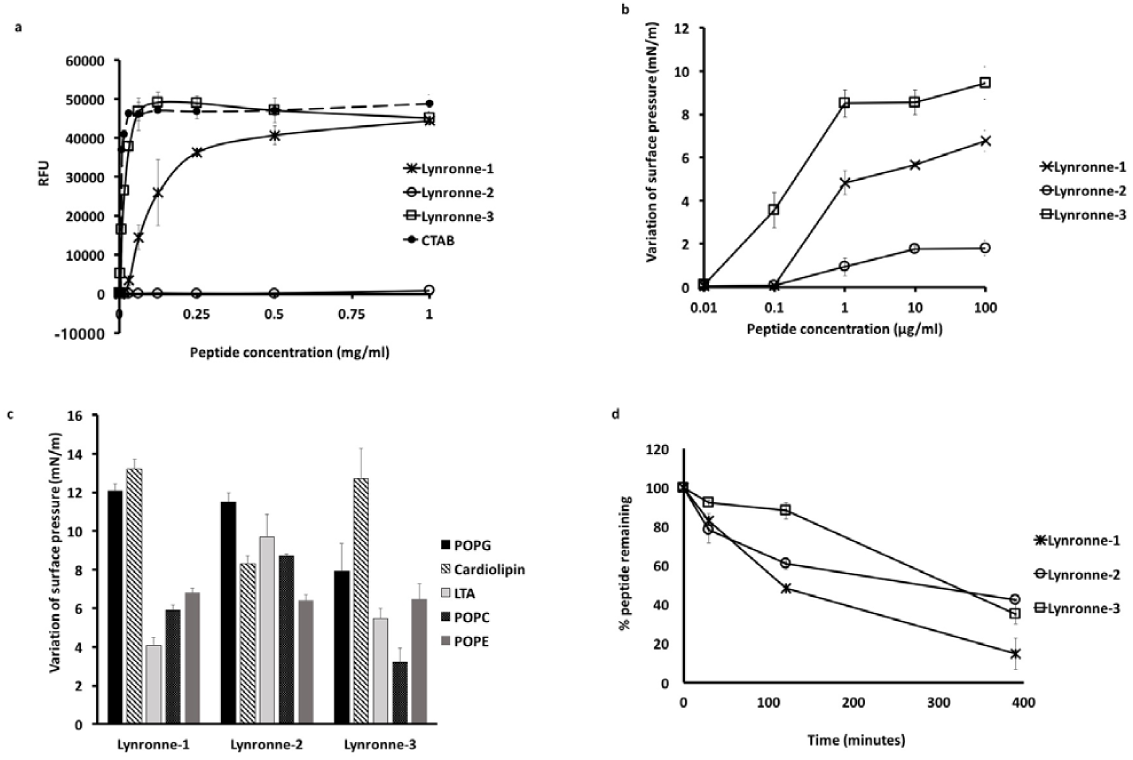
Mode of action studies of Lynronne-1, 2 and 3 against MRSA: **a)** Membrane permeabilization activity against MRSA USA300 measured by propidium iodide assay at 10 mins. **b)** Dose-dependent interaction of peptides with total MRSA lipid extract, the initial surface pressure of lipid monolayer being 30 + /-0.5 mN/m. **c**) Interaction of peptides (at 1 *μ* g/mL final concentration) with pure lipids-Maximal variation of surface pressure induced by the injection of peptide in lipid monolayer with initial surface pressure of 30 + /-0.5 mN/m. **d)** peptide stability/degradation in 25% serum over time. Relative peptide concentrations were determined by integration of the peaks from RP-HPLC chromatograms (values from three independent replicates; error bars represent one standard deviation).

Importantly, Lynronne-1, 2 and 3 also showed little haemolytic activity against red blood cells (Fig. 2d). Similarly, Lynronne-1, 2 and 3 were less cytotoxic to mammalian HUVEC and HepG2 cells when compared to the positive control agent, Staurosporine, which had 50% Growth Inhibition (GI_50_), Total Growth Inhibition (TGI) and 50% Lethal Concentration (LC_50_) of 0.00047, 0.0047 and 0.014 *μ* g/ml respectively. Lynronne-2 had no cytotoxic activity against mammalian HUVEC and HepG2 cells at the highest concentration tested (128 *μ* g/ml) while Lynronne-3 had little cytotoxicity at concentration close to the MIC with GI_50_, TGI and LC_50_ of 24.4, 61.2 and >128 *μ* g/ml respectively. Lynronne-1 on the other hand, had low toxicity at concentrations above the MIC with GI_50,_ TGI and LC_50_ of 45.9, 67.1 and 98.1 *μ* g/ml respectively. Whereas Lynronne-1 and 3 were able to permeabilize MRSA USA300 cell membrane, very little permeabilization activity was observed for Lynronne-2 even at >3 times its MIC concentration (Fig. 3a and Extended Data Fig. 4a), demonstrating that although antimicrobial activity of Lynronne-1 and 3 is due to pore formation, mechanism of action of Lynronne-2 may be different. In accordance with cytotoxicity results, Lynronne-1, 2 and 3 were less able to permeabilize eukaryotic membranes (Hep2G) compared to bacterial cell membranes (Extended Data Fig. 4b).

Since bacterial membrane permeabilization was demonstrated by the newly identified AMPs, we used lipid monolayer (Langmuir balance) to narrow down the potential lipids that are targeted by these peptides. The use of MRSA lipid extract demonstrated a higher insertion ability of Lynronne-1 and 3 compared to Lynronne-2 (Fig. 3b) and is in agreement with bacteria membrane permeabilization. The use of pure lipids demonstrated that Lynronne-1, 2 and 3 preferentially interact with lipids specific to bacteria, including 1-palmitoyl-2-oleoylsn-glycero-3-(phospho-rac-(1-glycerol)) (POPG), cardiolipin, lipoteichoic acid (LTA) or phosphatidylethanolamine (POPE) (also present in the inner membrane leaflet of eukaryotes) above lipids only present in eukaryotes, such as Palmitoyloleoylphosphatidylcholine (POPC) (Fig. 3c and Extended Data Fig. 4c, d). This may explain the limited haemolytic and cytotoxic activity against mammalian cells in spite of their antibacterial activity. Interestingly, Lynronne-2 showed the highest affinity for LTA demonstrated by measurement of the critical pressure of insertion of the peptides in the different lipids (Extended Data Fig. 4d), suggesting that this peptide could target other molecules containing teichoic acid moieties, including cell wall teichoic acids (WTA) and may also explain its antimicrobial activity in the absence of pore-formation in whole bacteria. Transmission electron micrographs of cells treated with Lynronne-1, 2 and 3 (at 3x MICs for 1 h) reveal varying changes in cell morphology and some cytoplasmic leakage (Fig. 4 and Extended Data Fig. 1b). MRSA USA300 cells treated with Lynronne-2 showed very little morphology changes, corresponding to the low permeabilization activity of the peptide. Further investigation into events leading up to changes in cell morphology may be required as the TEM analysis is semi-quantitative.

### *In vivo* efficacy

Given the promising *in vitro* activity of Lynronne-1, 2 and 3, we investigated their *in vivo* therapeutic potential within a murine model of MRSA skin infection^24^. Topical administrations of Lynronne-1 (2% w/v), in a group of mice with MRSA infected wounds (5 mice/group) once daily for three days was associated with a significant decrease (*P* < 0.01) in the colony count relative to the control group treated with phosphate buffered saline (PBS, pH 7.4) (Fig. 5a). However, this decrease was less than a 2-log reduction (≥99%) in colony counts. No significant decreases in colony counts were observed in the animal groups treated with Lynronne-2 (2% w/v) and Lynronne-3 (2% w/v). The reference, 2% mupirocin ointment (Dermasafe^®^ Ointment) was associated with a significant reduction (>2-log reduction, and *P* < 0.001) in bacterial counts, consistent with historical data ^24^. Given this promising *in vivo* result for Lynronne-1, we further tested its efficacy at a higher dose (10% w/v) in the wound model. Lynronne-1 at 10% (w/v) was associated with a significant (99%) reduction in bacterial density (≥2-log reduction, and *P* < 0.001) and comparable to mupirocin (2%) treatment (Fig. 5b). These findings suggest that Lynronne-1 can effectively reduce the bacterial load in MRSA infected wounds.

**Fig. 4.**
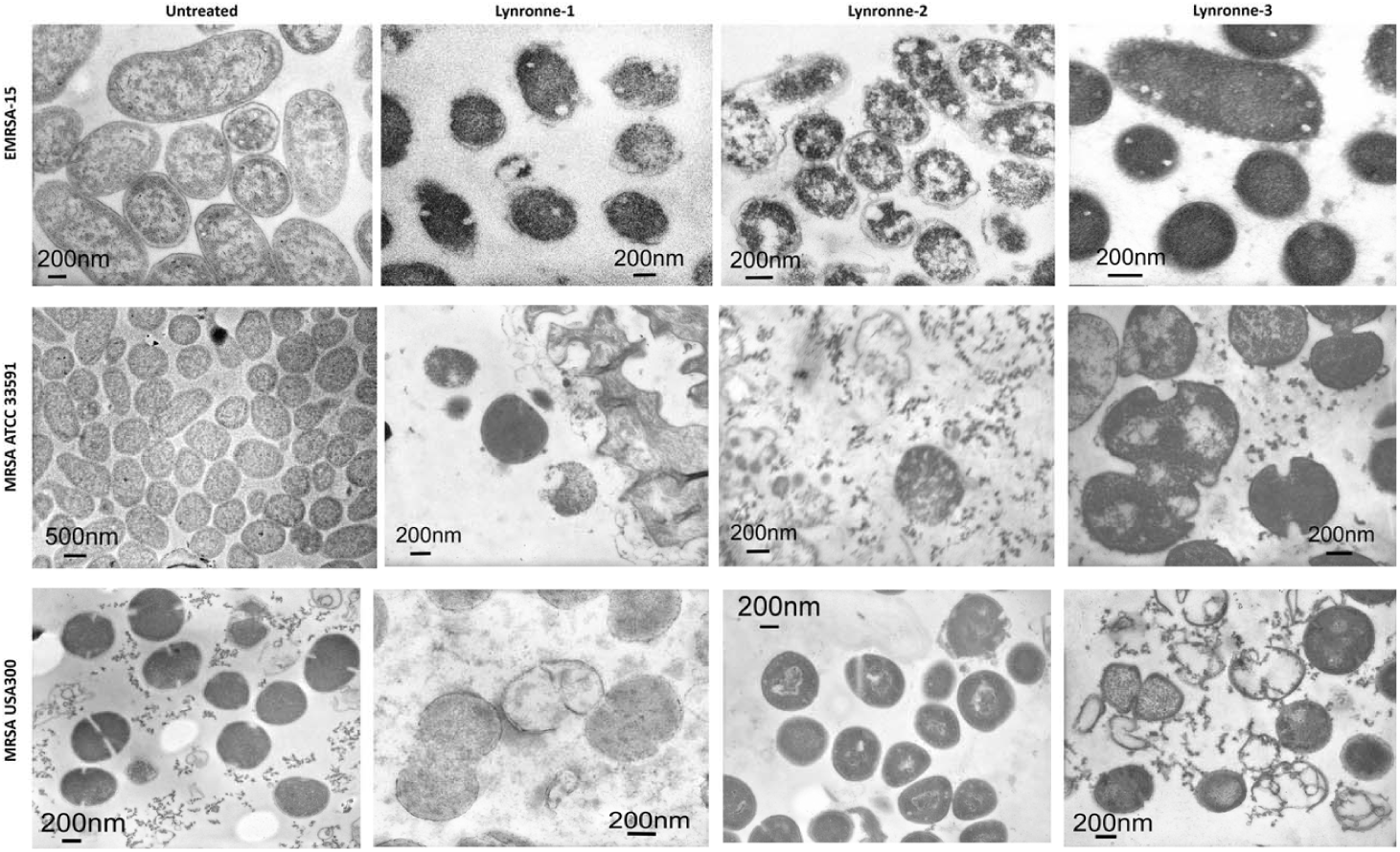
Transmission electron micrographs of MRSA cells. Representative micrographs of untreated and peptide treated MRSA cells. Cells were treated at 3x MIC peptide concentration for 1 h. Scale bars are shown on micrographs.

**Fig. 5.**
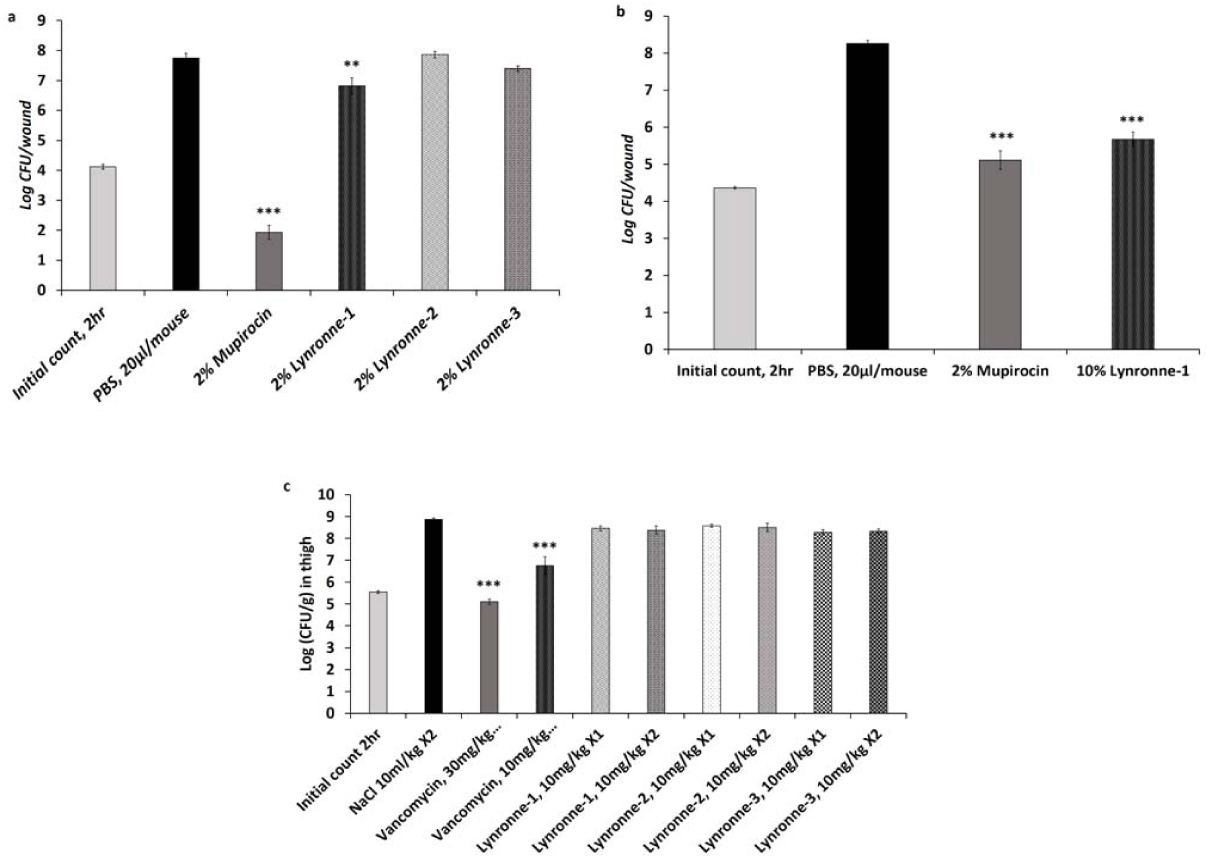
*In vivo* efficacy in mouse models of infection. **a)** Efficacy of Lynronne-1, Lynronne-2, Lynronne-3 and mupirocin against methicillin resistant *S.* USA300 MRSA (BAA-1717), wound infection in, ICR mice. Inoculum = 1.05 × 10^5^ CFU/ml. **b)** Efficacy of Lynronne-1 and mupirocin against methicillin resistant *S. aureus,* USA300 MRSA (BAA-1717). **c)** Efficacy of intravenously administered Lynronne-1, Lynronne-2, Lynronne-3 and Vancomycin in the *S. aureus* (ATCC 33591) MRSA thigh infection model with neutropenic male ICR mice. Inoculum = 1.57 × 10^5^ CFU/ml, x1 (administered once at 2 h) or x2 (twice at 2 and 8 h post infection). **\***, **\*\***, **\*\*\*)** Significant difference (*P* < 0.05, 0.01 and 0.001 respectively) compared to the control group (PBS or 0.9 % NaCl) was determined by one-way ANOVA followed by the use of a Dunnett’s test.

The efficacy of Lynronne-1, 2 and 3 were subsequently tested in a murine model of MRSA deep thigh infectiosn ^25^. The maximum tolerable dose (MTD) of all three AMPs was predetermined to be 10 mg/kg *in vivo* in mice. Test animals were inoculated intramuscularly (IM) in the right thigh with 0.1 ml/thigh of *S. aureus* MRSA ATCC 33591 (inoculum size of 1.57 × 10^5^ CFU/mouse) followed by treatment with Lynronne-1, 2 and 3 and the comparator antibiotic, vancomycin. Intravenous (IV) administrations of Lynronne-1, 2 and 3 at 10 mg/kg once or twice (2 h or 2 and 8 h post infection) were not associated with any significant antimicrobial effects compared to the vehicle/control group (0.9% NaCl). Vancomycin administered twice at 30 mg/kg IV) elicited a significant reduction (≥99%, 2-log reduction in CFU/g and P < 0.001). However, vancomycin at 10 mg/kg IV (administered twice) only elicited a significant reduction (P < 0.001) but not a 2-log reduction in colony counts compared to the control group (Fig. 5c).

Upon testing for stability of peptides, we observed that Lynronne-1 rapidly degraded in the presence of serum with only 48% and 15% of the parent peptide remaining after 2 h and 6 h respectively. Lynronne-2 and 3 were relatively more stable with ≥ 60% and 42%, as well as ≥88% and 34% after 2 h and 6 h respectively (Fig. 3d). A similar pattern was observed when these peptides were treated with trypsin (Extended Data Fig. 5), a common feature among peptides that are susceptible to degradation by peptidases^26^. This susceptibility to degradation may explain the poor efficacy of the peptides when administered intravenously.

## Discussion

Antimicrobial peptides are promising candidates for overcoming the critical and growing challenge of bacterial resistance to currently utilized antibiotics. The high cost and low success rate associated with the identification of novel and affective AMPs from natural sources has hindered the search for novel AMPs in the past. This hurdle has been overcome by new technological advances and improvements in traditional recombinant methods ^23^. Using a combination of functional metagenomics and computational approaches, we were able to show that the rumen microbiome is a promising resource for bio-prospecting novel AMPs, and these three peptides (Lynronne 1, 2 and 3), identified in this study are potential therapeutic candidates.

Lynronne 1, 2 and 3 are efficacious against clinically important drug-resistant pathogens in *in vitro* models of infection. In addition, Lynronne-1 decreased bacterial counts in MRSA wound infections using a murine model similar to commercially used mupirocin ointment, suggesting that it could be used topically in the treatment of MDR bacterial infections. Lynronne 1, 2 and 3 displayed low haemolytic activity against blood cells and negligible cytotoxicity against mammalian cells. Results presented here suggest that loss of cell viability after exposure to Lynronne-1, 2 and 3 among many factors, is due to membrane permeabilization, which contributes to membrane disruption and leakage of cell content. In addition to their broad antibacterial spectrum, selectivity and rapid killing of bacterial cells, Lynronne-1, 2 and 3 also showed a low tendency to select for resistance in the bacteria strains tested. The antimicrobial compounds discovered here are novel and demonstrate potent activity against clinically relevant human pathogens, rendering them as potential therapeutics. The identification of these novels AMPs support the hypothesis that the rumen is a promising resource for the discovery of novel antimicrobials with clinical relevance.

## Online Methods

### Functional antimicrobial activity screening of a rumen metagenomic library

A metagenomic library consisting of 8,448 clones from a plant-associated rumen microbiome was prepared previously ^15^. The clones were transferred gently using multichannel pin replicators (Molecular Devices Ltd, Berkshire UK) onto Luria-Bertani agar plates containing a lawn (500 *μ* l) of *Staphylococcus aureus* (MSSA) RN4220, *Escherichia coli* K12, *Salmonella enterica* serovar Typhimurium SL1344, *Enterococcus faecalis* JH2-2 and *Listeria monocytogenes* NCTC 11994 serovar 4b (OD_600_ = 1) using sterile pin replicators. Plates containing these pathogens were incubated at 37^°^C (or 30^°^C for *L. monocytogenes*) for 24 h, and clearing zones around the fosmid clones were considered as indicative of clones containing gene inserts coding for antimicrobial compounds. Assays were performed in sextuplicates to ensure reproducibility.

### Sequence-based screening of putative antimicrobial positive fosmid clones

Fosmids with putative antimicrobial activity were sequenced using the Genome Sequencer FLX platform (454 Life Sciences) according to manufacturer guidelines. Following sequence assembly using the default parameters on the gsAssembler software v.2.5.3 (Roche Life Sciences), sequences were trimmed and vector sequence were removed using BioEdit (version 7.1.11) ^27^. VecScreen on NCBI was used to search the sequences for vector contamination. Open reading frames (ORFs) were determined using NCBI’s ORF finder on the NCBI blast suite (v2.2.28) ^28^.

### *In silico* identification of antimicrobial peptides

Prediction of antimicrobial peptide sequences within the identified ORFs was achieved using a variety of software available at various public databases. Specifically, we used the antimicrobial peptide database (APD) ^29^, the Antimicrobial Sequence Scanning System (AMPA) ^30^, BACTIBASE ^31^, Collection of Anti-Microbial Peptides (CAMP) ^18,32^, and Antibacterial peptides (AntiBP) ^33^. Only ORFs that were predicted to encode putative antimicrobial peptides within at least two of the utilized databases were explored further. The active sites within positive ORFs were predicted using AMPA ^30^ and make up the peptides included in the peptide library to be screened for antimicrobial activity.

### Peptide synthesis and antimicrobial screening

The peptides in the created peptide library were synthesized based on the SPOT technique and pure peptides (≥95% purity) were synthesized on resin using solid phase Fmoc peptide chemistry ^16^. The 135 peptides in the peptide library was screened for antimicrobial activity using both luminescent (*Pseudomonas aeruginosa* strain H1001) ^34^ and non-luminescent (epidemic methicillin resistant *S. aureus* EMRSA-15, *Sal. typhimurium* and *E. coli* K12) bacterial strains. Activity against non-luminescent bacteria strains was assessed by fluorescence readings (excitation/emission spectra of 560/590 nm) after the addition of resazurin dye (100 *μ* m final concentration). The results from the screen were analysed using the MATLAB extension Gait-CAD as previously described ^35^. Concentration values with a 75% reduction of luminescence or fluorescence (IC75) in relation to untreated controls were computed. Three peptide activity classes: active (IC75≤0.25), semi-active (0.25<IC75≤0.50) and inactive IC75>0.5 were assigned against all microorganisms.

### Determination of minimum inhibitory concentration (MIC)

Minimum inhibitory concentrations (MICs) were determined using a modified broth microdilution method ^36^ in cation adjusted Mueller Hinton broth (MHB) following the International Organization for Standardization (ISO) 20776-1 standard for MIC testing with a final bacterial inoculum concentration of 5 × 10^5^ CFU/ml ^37^. Peptides dissolved in sterile distilled water and comparator antibiotics dissolved in the appropriate solvents ^38^ were added to sterile U-bottom polypropylene 96 well microplates at desired concentrations. MIC was defined as the lowest concentration of peptide or antibiotic, which inhibited the visible growth of bacteria after 18-24 h incubation at appropriate temperatures.

### Time kill kinetics

Assessment of the bactericidal activity of peptides and comparator antibiotics was performed as previously described ^39^ using exponential-phase cultures of pathogens grown in MHB (1 × 10^8-10^ CFU/ml). Peptides were added at concentrations 3 times their MIC values. Experiments were performed in quadruplicates, and CFU/ml was calculated at different time points after over overnight incubation.

### Selection for AMP resistance

To evaluate if populations of AMP-resistant bacteria could be selected, cultures were continuously exposed to Lynronne-1, 2 and 3 for a duration of 25 days, as previously described ^40^. Briefly, broth microdilution susceptibility testing was performed using a standard doubling-dilution series of Lynronne-1, 2 and 3 concentrations on day 1. Following incubation of the cultures for 24 h, and determination of the MIC, the well that contained the highest concentration of AMPs permitting growth was diluted 1: 1000 in MHB and used to provide the inoculum for the next MIC assay; this process was repeated daily for 25 days.

### Erythrocyte leakage assay

The haemolytic activity of peptides was determined by measuring haemoglobin leakage from human and sheep erythrocytes (Divbioscience, NL) after 1 h incubation at OD_450_nm. Cetyltrimethylammonium bromide (CTAB), a non-specific detergent was used also included as a positive control. Triton-X 100 (0.1% v/v) was also used as an indicator of 100% haemolysis while PBS treatment represented auto-haemolysis.

### Cytotoxicity of peptides

Cytotoxic activity of the identified peptides was determined using HUVEC and HEPG2 cells as described previously ^41,42^. Three dose response parameters 50% Growth Inhibition (GI_50_), Total Growth Inhibition (TGI) and 50% Lethal Concentration (LC_50_) were calculated for each experimental agent and cellular tumor assay results are presented as the percent cell growth or percent tube length of specific activity. Significant responses were defined when observed cell growth was ≤ 50% and tube length was ≤ 70% using the proliferation and tube formation assays respectively. Permeabilization of eukaryotic membrane by Lynronne-1, 2 and 3 was also investigated using HepG2 cells as described previously ^43^.

### Membrane permeabilization and peptide-lipid interactions

Membrane permeabilsation was evaluated using propidium iodide assay as previously explained ^43^, with CTAB as positive control. Peptide-lipid interaction was measured using reconstituted lipid monolayer ^44^. Total lipid extract from overnight liquid cultures of MRSA USA300 was obtained by Folch extraction, resuspended in chloroform and stored at −20°C under nitrogen conditions. Pure bacterial and eukaryotic lipids, POPG, cardiolipin, LTA, POPE and POPC (Avanti Polar Lipid USA) were reconstituted in chloroform at 1 mg/ml and stored at-20°C under nitrogen. Using a 50 *μ* l Hamilton’s syringe, total MRSA lipid extract or pure lipids were spread at the surface of PBS creating a lipid monolayer at the air-water interface until an initial surface pressure of 30 + /-0.5 mN/m was reached. This corresponds to a lipid packing density theoretically equivalent to that of the outer leaflet of the cell membrane ^45^. After 5-10 mins of incubation allowing evaporation of the solvent and stabilization of the initial surface pressure, peptides were injected into the PBS (pH 7.4, volume 800 *μ* l) sub-phase using a 10 *μ* l Hamilton syringe. The variation of the surface pressure caused by peptide injection was then continuously monitored using a fully automated microtensiometer (μTROUGH SX, Kibron Inc., Helsinki, Finland) until reaching equilibrium (maximal surface pressure increase usually obtained after 15 to 25 mins). Critical pressure of insertion of each peptide in the different lipids was also determined by changing the initial pressure of lipid monolayer (from 10 and 30 mN/m) and measuring the variation of pressure caused by the injection of peptide (at 1 *μ* g/ml final concentration). All experiments were carried out in a controlled atmosphere at 20^°^C ±1^°^C and data were analyzed using the Filmware 2.5 program (Kibron Inc., Helsinki, Finland). Variation of surface pressure was plotted as a function of initial surface pressure and critical pressure of insertion was calculated as the theoretical value of initial pressure of lipid monolayer not permissive to peptide insertion, i.e. a variation of pressure equal to 0 mN/m. The accuracy of the system under our experimental conditions was determined to be ±0.25 mNm/1 for surface pressure measurements.

### Serum stability and trypsin degradation assays

The stability of Lynronne-1, 2 and 3 in the presence of serum was investigated by monitoring the concentrations of the remaining soluble peptides by reverse phase high performance liquid chromatography (RP-HPLC), as described previously by Nguyen and colleagues ^26^. Degradation of Lynronne-1, 2 and 3 in the presence of trypsin was adapted from a previously described method ^46^. Briefly, trypsin (5 *μ* l of 0.5 ng/ml) and 37.5 *μ* l trypsin activation buffer (50 mM Tris, 2 mM CaCl_2_, pH 7.8-8) was added to 7.5 *μ* l of 5 mg/ml Lynronne-1, 2 and 3 and incubated at 37 °C for different time points (0, 1, 3, and 24 h). An aliquot from each reaction mixture at 0 h was prepared to allow matrix-assisted laser desorption/ionization time-of-flight (MALDI-TOF) mass spectrometric verification of the synthetic peptides prior to incubation. Trypsin action buffer (5 *μ* l) and peptides without trypsin was used as negative control. After the various incubation times, the reaction mixtures were acidified by addition of an equal volume (50 *μ* l) of 10 % (v/v) trifluroacetic acid (TFA) to stop further enzyme activity. Lynronne-1, 2 and 3 degradation products were evaporated to dryness and reconstituted in acetonitrile/water/TFA (40/59.5/0.5 %, v/v/v). Samples (1 *μ* l) were carefully placed onto a stainless steel (MALDI) target, covered with 1 *μ* l of matrix (53 mM a-cyano-4-hydroxycinnamic acid in acetonitrile/water/TFA, 70/29.97/0.03 %, v/v/v) solution before analysis by mass spectrometry.

### Molecular modelling of antimicrobial peptide 3D structures

The modeling of the 3D conformation of peptides was carried out using the *de novo* structural prediction method PEP-FOLD ^17^. The best 3D models for each peptide was selected according to PEP-FOLD score, based on the OPEP force field ^47^. Results were visualized using the PyMOL v1.7.6 program ^48^.

### Transmission electron microscopy

Effects of AMPs on bacterial cell morphology was investigated using transmission electron microscopy as described by Huws *et al* ^49^. Bacterial cultures in mid-log phase were treated with Lynronne-1, 2 and 3 (at 3x MIC for 1 h) and then fixed with 2.5% (v/v) glutaraldehyde. Cells were post-fixed with 1% osmium tetroxide (w/v), stained with 2% (w/v) uranyl acetate and Reynold’s lead citrate and observed using a JEOL JEM1010 transmission electron microscope (JEOL Ltd, Tokyo, Japan) at 80 kV.

### Quantification of anti-biofilm activity

The effectiveness of Lynronne-1, 2, and 3 to prevent a) biofilm attachment and b) disrupt established biofilms was measured using a 96 well biofilm model ^50^. Briefly, MRSA cultures grown in Brain Heart Infusion (BHI) broth were washed, resuspended to an OD600nm = 0.02 in BHI broth and a) treated with peptides (at 1/4x, 1/2x, 1x, 2x and 4x MIC) and incubated at 37 °C for 24 hours or b) incubated at 37 °C for 24 h to allow establishment of biofilms, washed in PBS to remove planktonic cells, and further incubated with peptides (at 1/4x, 1/2x, 1x, 2x and 4x MIC) in fresh BHI broth at 37°C for 24 h. The biofilms were washed in PBS to remove planktonic cells, fixed with methanol, stained with 0.5% (w/v) crystal violet and re-solubilized with 33% (v/v) acetic acid. The optical density of biofilms was measured at 570_nm_.

### *In vivo* efficacy of peptides in a mouse model of MRSA skin and thigh infections

The efficacy of Lynronne-1, 2 and 3 for treating a wound infection caused by USA300 community-associated methicillin resistant *S. aureus* (MRSA, strain BAA-1717) was evaluated using a murine wound model with ICR mice ^24^. A sharp punch (ID 12 mm) was used to create excisional skin wounds in groups of five female ICR mice (5-6 weeks old). The mice were inoculated topically on the wound area (5 *μ* l per wound) with 1.05 × 10^5^ CFU/mouse of strain BAA-1717 suspended in 5 *μ* l PBS. Peptides (2% or 10% w/v) in PBS (pH 7.4), control (PBS), and mupirocin 2% (Dermasafe^®^ ointment) were topically applied (20 *μ* l/mouse) directly to wounds 2 h post infection and once daily thereafter for 3 consecutive days. Test animals were sacrificed and the wound tissue was excised for enumeration of bacteria cells (CFU/wound). A two-log reduction in the bacterial count (≥99% reduction) relative to the control group indicates significant activity. Statistical significance (*P* < 0.05) was performed with one-way ANOVA followed by use of the Dunnett’s method.

In the thigh infection model, using neutropenic mice ^25^, groups of five male ICR mice weighing 22 ± 2 g were inoculated (1.57 × 10^5^ CFU/mouse (0.1 ml/thigh)) intramuscularly (IM) with *S. aureus* (MRSA, ATCC 33591) in the right thigh. Lynronne-1, 2 and 3 at 10 mg/kg were intravenously (IV) administered once 2 h post-infection or twice 2 and 8 h post-infection. The reference substance, vancomycin, was also intravenously injected 2 and 8 h post-infection at 10 and 30 mg/kg. Test animals were sacrificed 26 h after inoculation and the infected thighs were excised and the bacterial burden (CFU/gram) was determined.

## Data availability statement

The datasets generated and/or analysed during the current study are available under the study name [SRP099633 Cow rumen Metagenome] in the GenBank database, [https://www.ncbi.nlm.nih.gov/].

## Acknowledgements

This project was funded partly by the Cross River State Government of Nigeria, the Life Sciences Research Network Wales, RCUK Newton Institutional Link Fund (172629373), and the BBSRC UK (BB/L026716/1). PNG acknowledges ERA NET IB2 (grant Nr ERA-IB-14-030) funded through the BBSRC grant BB/M029085/1. We are also grateful to Teri Davies for her technical assistance in setting up some experiments and to Dr Colin Greengrass, Prof Andrea Brancale and Prof Keith Harding for their advice. The *in vivo* mouse studies were carried out by Eurofins Panlabs.

## Author Contributions

LO and SH conceived the project. LO, with help from HV, TW and MW, completed the laboratory work under supervision of SH, JE and CC. SG and LO completed the sequencing and downstream analysis of the sequences respectively. AC and NF assisted LO with transmission electron microscopy and 3D structural modelling respectively. FP, OG and PG created the rumen fosmid metagenome library. LO, MM and JP completed the membrane permeabilization and lipid binding assays. LO, YL, FL and MZ completed the serum stability and degradation assays. KH and RM assisted LO with peptide library creation, screening and data analysis. MH and HM have provided valuable ideas into the project from the time of conception. LO wrote the paper with input from all co-authors.

## Competing financial interests

There are no conflicting financial interests to declare.

Correspondence and requests for materials should be addressed to.

## Extended Data Tables

**Extended Data Table 1:** Assembly metrics of selected clones after shotgun sequencing using the Roche GS FLX system.

| Fosmid clone ID | GenBank BioSample ID | Total number of reads | Total number of base pairs | Number of contigs | Total length of contigs (bp) | Insert size from chosen contig (bp) | GC content of insert (%) | Number of ORFs in contig |
| --- | --- | --- | --- | --- | --- | --- | --- | --- |
| SAB PL4 N2 | SAMN063330279 | 26413 | 17318197 | 1102 | 2102927 | 25841 | 56.2 | 132 |
| SAB PL5 A1 | SAMN063330280 | 47599 | 30986482 | 691 | 4325733 | 67719 | 52.7 | 328 |
| SAB PL5 A2 | SAMN063330281 | 22427 | 14871695 | 1218 | 2450352 | 38145 | 51.5 | 194 |
| SAB PL5 C17 | SAMN063330282 | 5733 | 3699135 | 76 | 131887 | 12070 | 52.1 | 53 |
| SAB PL12(1) H18 | SAMN063330283 | 10168 | 6616004 | 203 | 263016 | 401 | 43.8 | 6 |
| SAB PL8 L14 | SAMN063330284 | 42326 | 26689339 | 1258 | 3662870 | 8036 | 47.9 | 45 |
| SAB PL9 A12 | SAMN063330285 | 4707 | 3113068 | 85 | 90709 | 2240 | 53.6 | 11 |
| SAB PL9 B3 | SAMN063330286 | 15899 | 10345946 | 387 | 572057 | 31665 | 48.2 | 160 |
| SAB PL9 F16 | SAMN063330287 | 10035 | 6672761 | 302 | 448066 | 13987 | 51.9 | 84 |
| SAB PL12(1) C3 | SAMN063330288 | 6626 | 4307213 | 82 | 121460 | 27728 | 52.6 | 149 |
| SAB PL12(1) D18 | SAMN063330289 | 10742 | 6936903 | 284 | 392503 | 8036 | 47.9 | 44 |
| SAB PL12(2) A3 | SAMN063330290 | 32909 | 21294400 | 1261 | 3447744 | 27125 | 47.7 | 168 |
| SAB PL12(2) G13 | SAMN063330291 | 17172 | 10597944 | 885 | 1449480 | 16816 | 51.1 | 34 |
| SAB PL12(2) J19 | SAMN063330292 | 26621 | 16046320 | 1290 | 2782997 | 31708 | 49.4 | 39 |
| SAB PL19 A3 | SAMN063330293 | 18860 | 11251168 | 645 | 992753 | 5795 | 53.4 | 33 |
| SAB PL27 G15 | SAMN063330294 | 22181 | 13408286 | 890 | 1518463 | 29772 | 45.5 | 61 |
| SAB PL27 G18 | SAMN063330295 | 14752 | 9075890 | 552 | 817667 | 24482 | 39.5 | 99 |
| SAB PL27 H14 | SAMN063330296 | 20027 | 12073019 | 950 | 1581893 | 12018 | 50.2 | 68 |
| SAB PL27 L10 | SAMN063330297 | 14117 | 8650852 | 349 | 514455 | 36694 | 51.0 | 193 |
| SAB PL27 L13 | SAMN063330298 | 14336 | 8770311 | 353 | 544607 | 35970 | 52.6 | 182 |
| SAB PL27 L17 | SAMN063330299 | 17428 | 10680537 | 751 | 1223750 | 25007 | 47.5 | 124 |
| SAB PL29 H11 | SAMN063330300 | 16143 | 10004756 | 648 | 988569 | 33998 | 49.3 | 176 |
| SAB PL29 I11 | SAMN063330301 | 21218 | 12829662 | 1101 | 1996994 | 23851 | 43.6 | 118 |
| SAB PL29 M21 | SAMN063330302 | 13816 | 8488354 | 581 | 842746 | 1609 | 56.6 | 7 |

**Extended Data Table 2:**
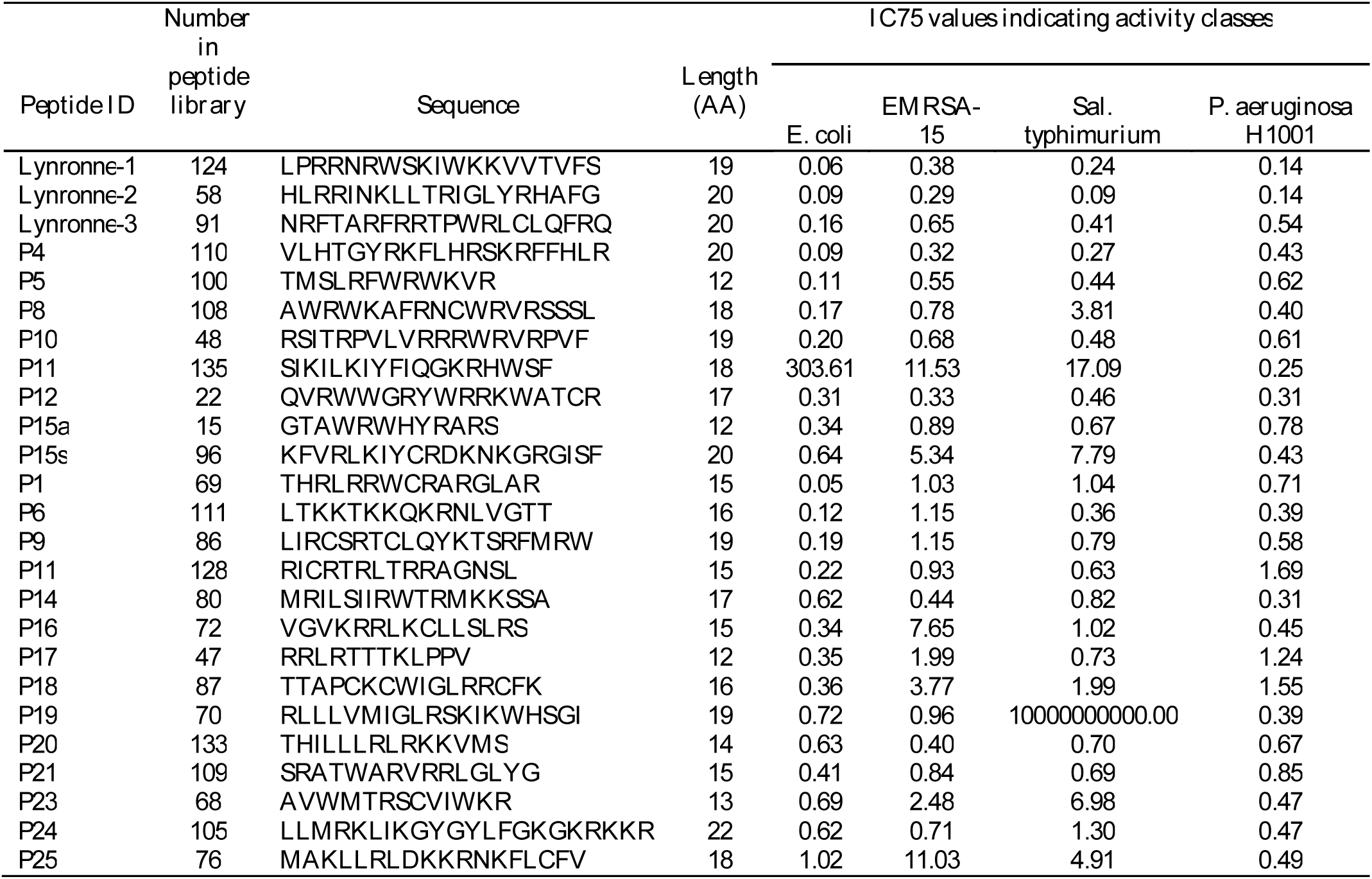
All active peptides in peptide library screen. All 25 peptides in peptide library with ≥75% reduction of luminescence/fluorescence (IC75) in relation to untreated controls. Three activity classes: active (IC75≤0.25), semi-active (0.25<IC75≤0.50) and inactive IC75>0.5.

### Extended Data Figures

**Extended Data Figure 1:**
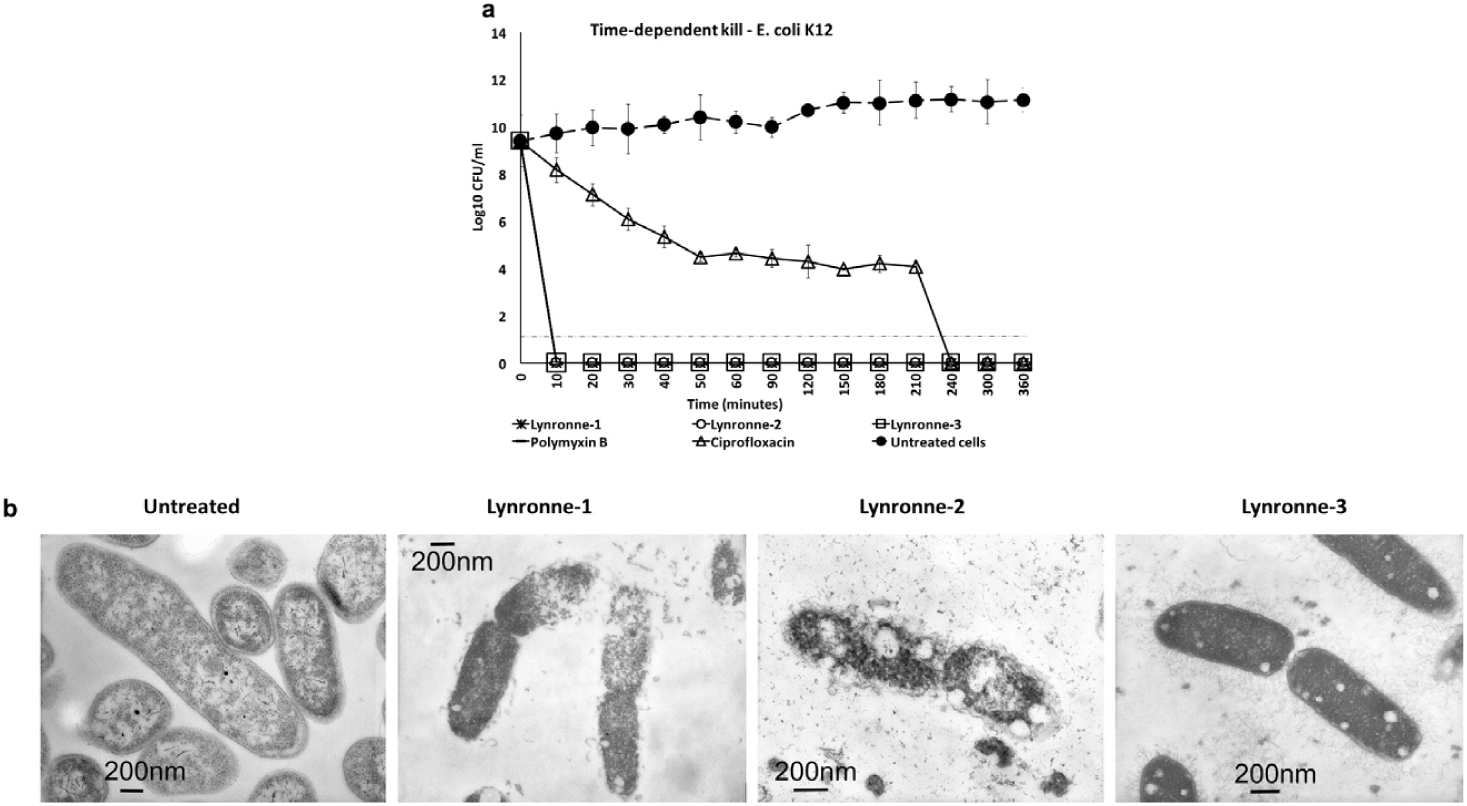
Time dependent kill of *E. coli* K12 by antimicrobial compounds (values from four independent replicates; error bars represent the standard deviation), dashed line indicates limit of detection **b)** Representative TEM images of *E. coli* K12 following 60-minute exposure to peptides at 3x MIC concentrations. Scale bars are shown on micrographs.

**Extended Data Figure 2:**
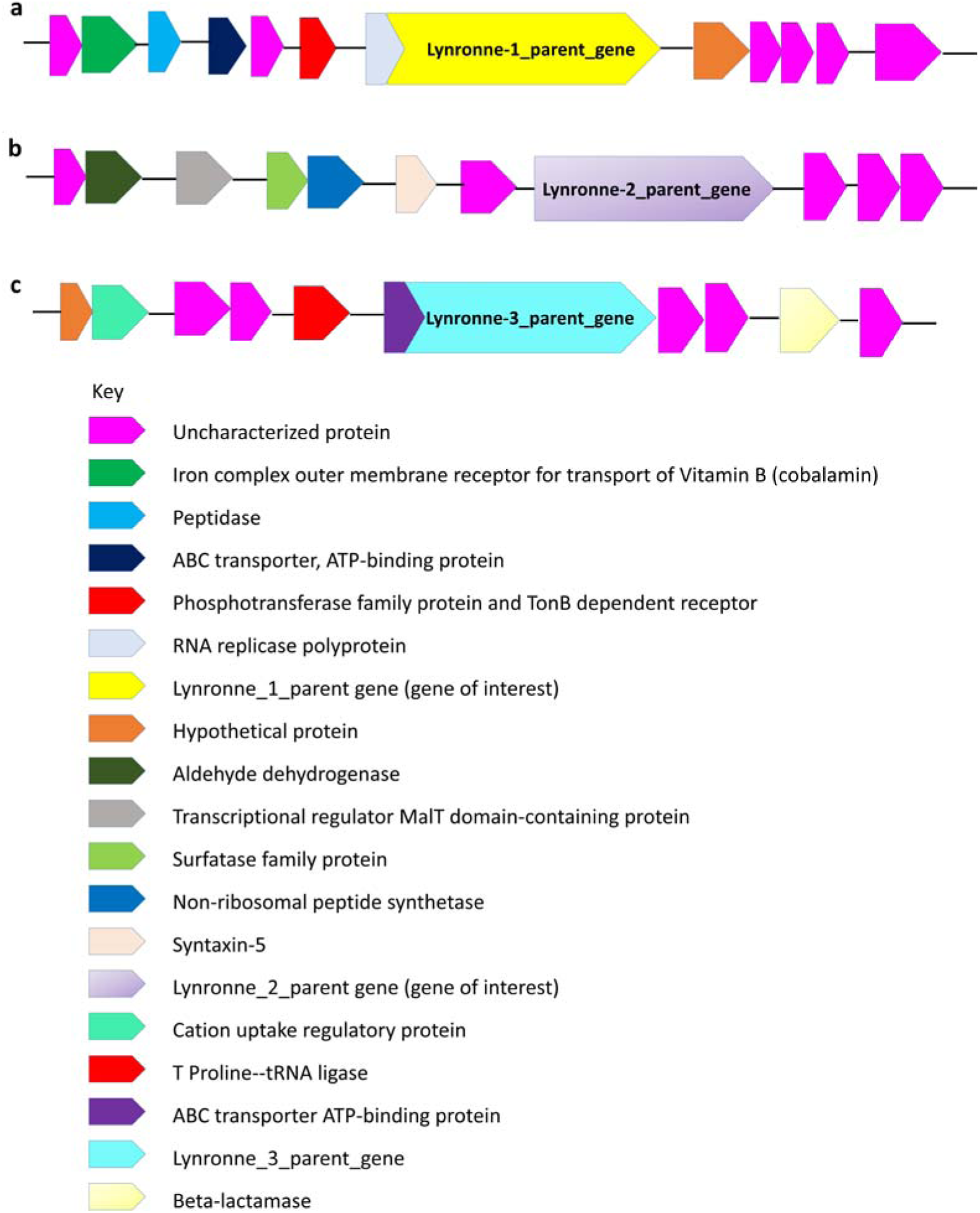
Sequences upstream and downstream of parent gene for. **a)** Lynronne-1, **b)** Lynronne-2 and **c)** Lynronne-3 predicted by NCBI blastp searches.

**Extended Data Figure 3:**
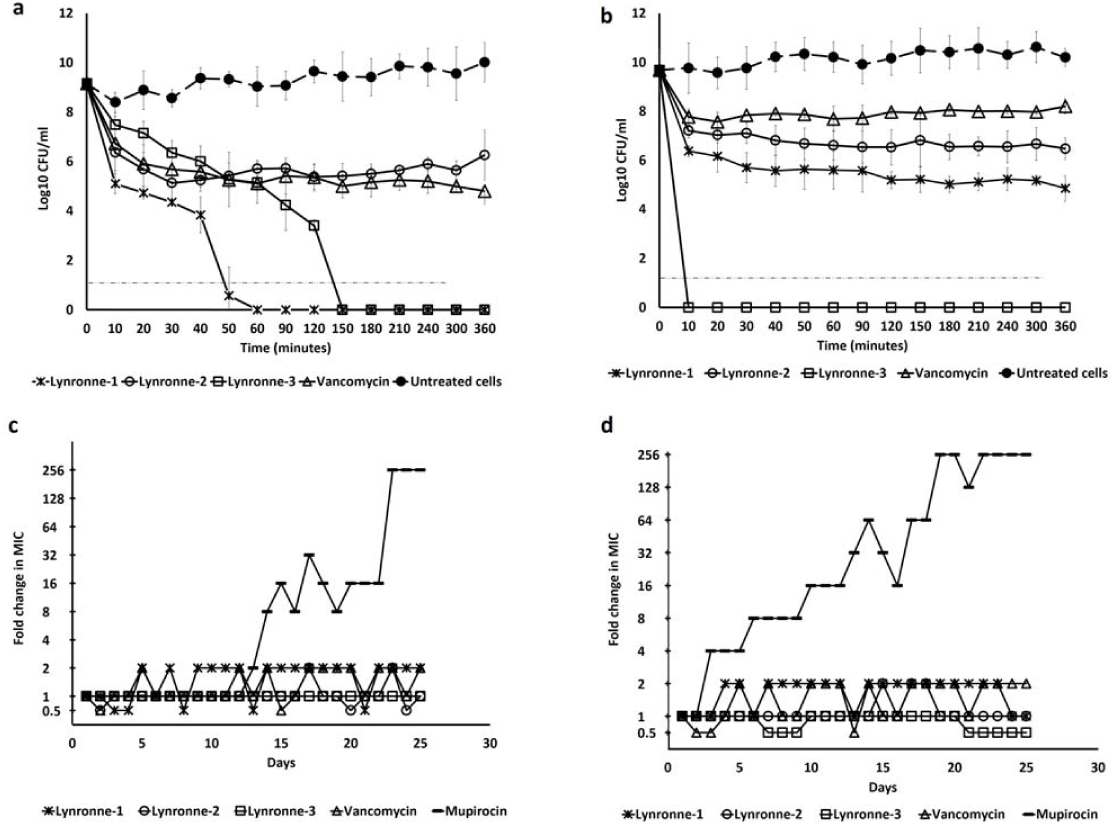
Antimicrobial activity of peptides: **a)** Time dependent kill of EMRSA-15 **b)** Time dependent kill of MRSA ATCC 33591 by antimicrobial compounds at 3x MIC concentration. Dashed lines indicate limit of detection. Values from three independent replicates; error bars represent one standard deviation. Resistance acquisition during serial passaging of **c)** EMRSA-15 and **d)** MRSA ATCC 33591 cells in the presence of sub-MIC levels of antimicrobials. The y axis is the fold change in MIC during passaging. For mupirocin, 256x MIC was the highest concentration tested. The figure is representative of 3 independent experiments.

**Extended Data Figure 4:**
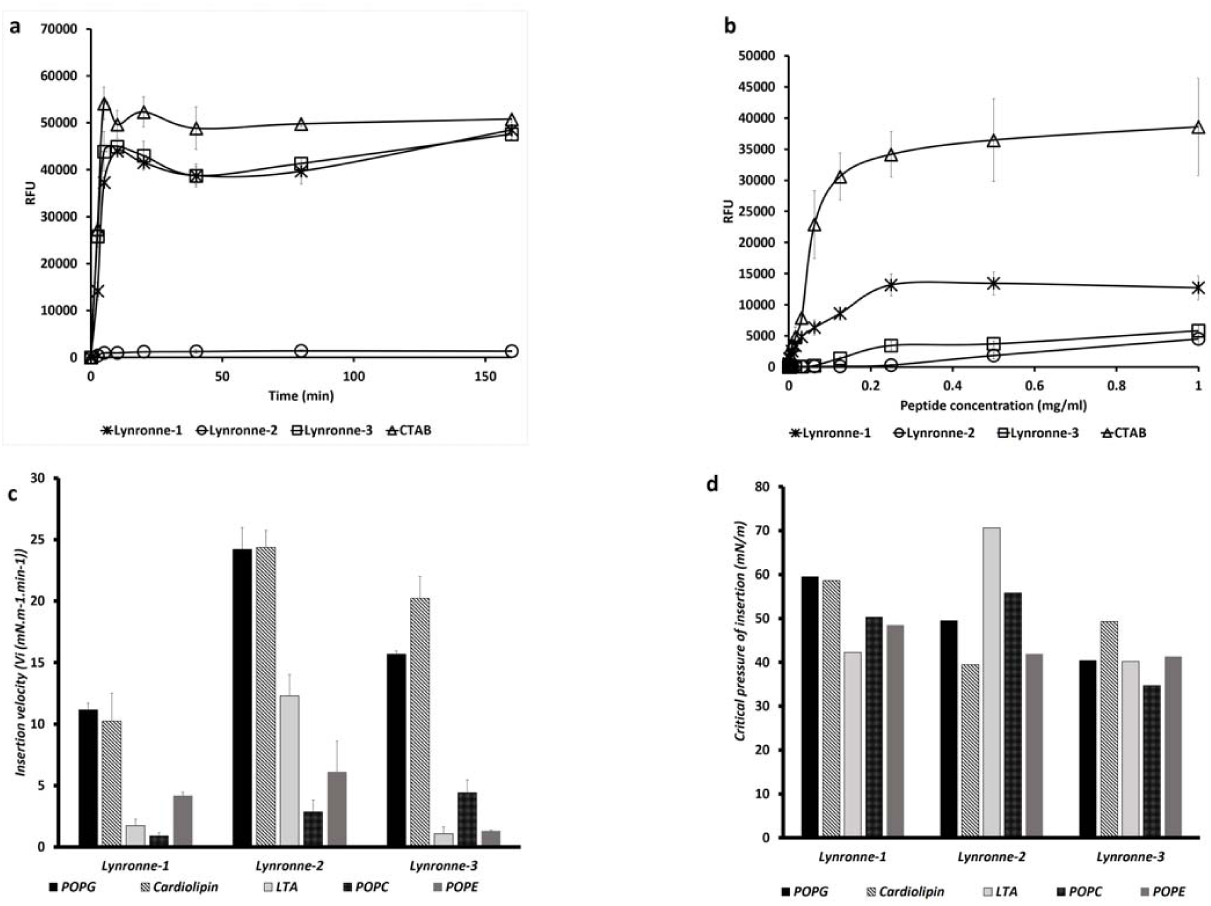
Mode of action of peptides: **a)** Time dependent membrane permeabilzation of MRSA USA 300 cells by peptides at 1 mg/ml. **b)** Dose-effect of peptides on HepG2 permeabilization measured by propidium iodide assay at 10 mins. **c)** Interaction of peptides (at 1 *μ* g/mL final concentration) with pure lipids-Insertion velocity into pure lipids and **d)** Critical pressure of insertion (mN/m) of Lynronne-1, 2 and 3 obtained by measuring the variation of surface pressure induced by the injection of peptide in lipid monolayer with various initial surface pressures.

**Extended Data Figure 5:**
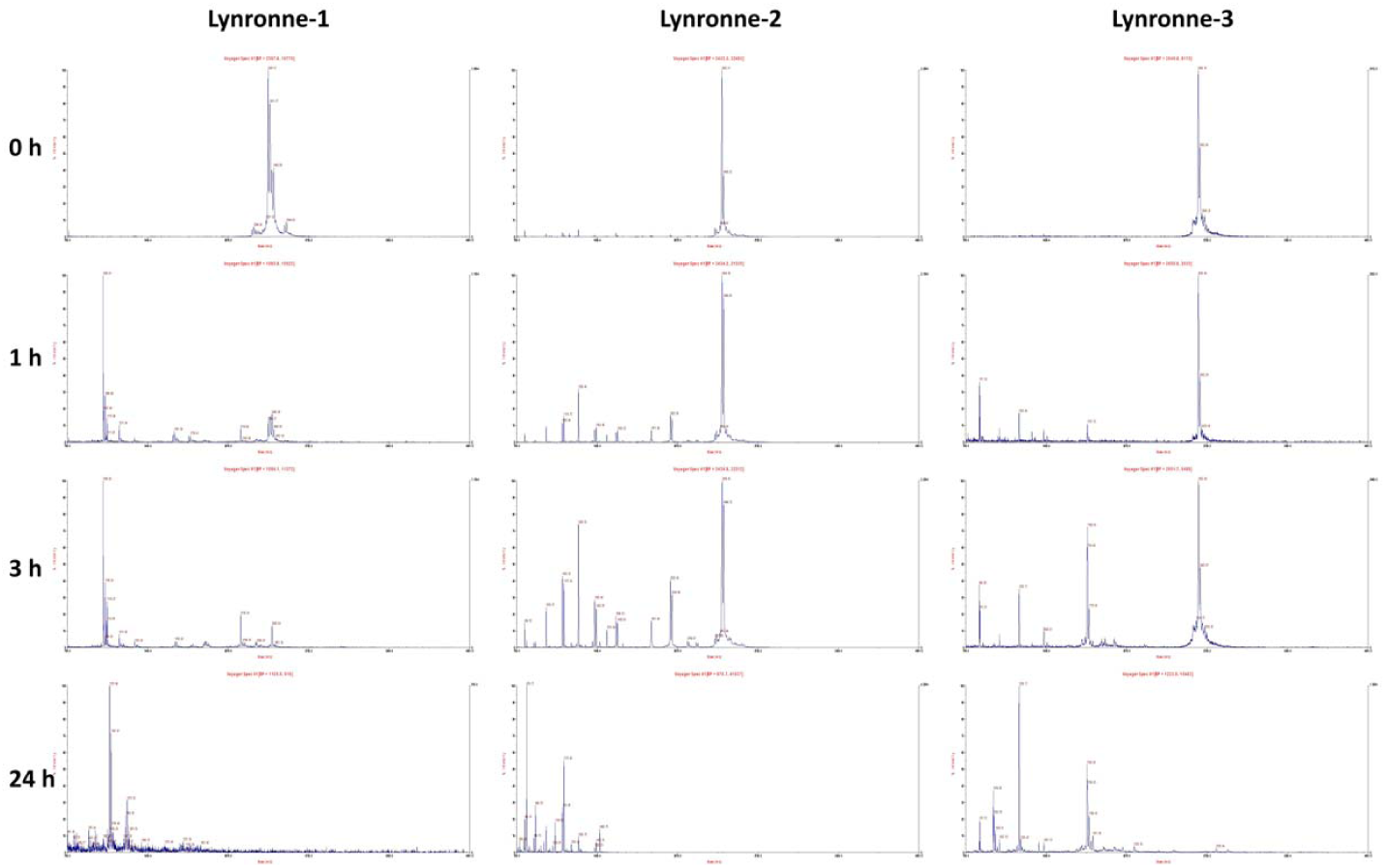
Peptide stability/degradation in the presence of trypsin. An example set of MALDI-TOF chromatograms from various time points of peptides incubated with trypsin (final concentration of 0.05 ng/ml) at 37°C, showing the non-degradation/degradation of the intact peptide and appearance of its partially digested products.

